# Reduced Auditory Steady State Responses in Autism Spectrum Disorder

**DOI:** 10.1101/837633

**Authors:** R.A Seymour, G Rippon, G Gooding-Williams, P.F. Sowman, K Kessler

**Affiliations:** Aston Neuroscience Institute, School of Life and Health Sciences, Aston University, Birmingham, B4 7ET; Department of Cognitive Science, Macquarie University, Sydney, Australia, 2109; Wellcome Centre for Human Neuroimaging, Queen Square Institute of Neurology, University College London, 12 Queen Square, London WC1N 3AR, UK

**Keywords:** Auditory Steady State, Autism Spectrum Disorder, MEG, Gamma, Sensory

## Abstract

**Background:** Auditory steady state responses (ASSRs) are elicited by clicktrains or amplitude-modulated tones, which entrain auditory cortex at their specific modulation rate. Previous research has reported reductions in ASSRs at 40Hz for autism spectrum disorder (ASD) participants and first-degree relatives of people diagnosed with ASD [1,2].

**Methods:** Using a 1.5s-long auditory clicktrain stimulus, designed to elicit an ASSR at 40Hz, this study attempted to replicate and extend these findings. Magnetencephalography (MEG) data were collected from 18 adolescent ASD participants and 18 typically developing controls.

**Results:** The ASSR localised to bilateral primary auditory regions. Regions of interest were thus defined in left and right primary auditory cortex (A1). While the transient gamma-band response (tGBR) from 0-0.1s following presentation of the clicktrain stimulus was not different between groups, for either left or right A1, the ASD group had reduced oscillatory power at 40Hz from 0.5 to 1.5s post-stimulus onset, for both left and right A1. Additionally, the ASD group had reduced inter-trial coherence (phase consistency over trials) at 40Hz from 0.64-0.82s for right A1 and 1.04-1.22s for left A1.

**Limitations:** In this study, we did not conduct a clinical autism assessment (e.g. the ADOS), and therefore it remains unclear whether ASSR power and/or ITC are associated with the clinical symptoms of ASD.

**Conclusion:** Overall, our results support a specific reduction in ASSR oscillatory power and inter-trial coherence in ASD, rather than a generalised deficit in gamma-band responses. We argue that this could reflect a developmentally relevant reduction in non-linear neural processing.

## Background

Autism Spectrum Disorder (ASD) is a neurodevelopmental condition characterised by impairments in social interaction, disrupted communication and repetitive behaviours [3]. Although these features remain the primary diagnostic markers of ASD, the presence of sensory symptoms has recently been given a more central diagnostic role. This change in symptom emphasis reflects the observation that over 90% of ASD individuals experience hyper- and/or hypo-sensitive sensory perception [4,5]. It has been suggested that differences in low-level sensory processing contribute to the atypical developmental trajectories of higher-level cognitive functions in autism [6]. An understanding of the neural circuits involved will therefore prove fruitful for ASD research, and could even facilitate the identification of earlier, brain-based diagnostic markers [7,8].

Dysregulated neural oscillations are a promising neural correlate of atypical sensory processing in ASD. In particular, atypicalities in high frequency gamma-band oscillations (30-80Hz) have been reported in ASD across visual, auditory and somatosensory domains [7,9–13]. Gamma oscillations are generated through excitatory-inhibitory (E-I) neuronal coupling [14], which facilitates periods of pre- and post-synaptic excitability alignment, thereby promoting efficient neural communication [15]. Findings of atypical gamma oscillations in ASD may therefore reflect disrupted E-I interactions within cortical micro-circuits [16], and concomitant effects on local and global brain connectivity [17].

Within the context of auditory processing, dysregulated gamma-band oscillations in ASD have been previously reported [7]. One prevalent approach to study auditory gamma-band activity non-invasively is through amplitude modulated tones called “clicktrains”. Such stimuli produce two distinct gamma-band responses. First, a transient gamma-band response (tGBR) is generated within 1s of stimulus onset [18]. This tGBR spans frequencies from 30-60Hz and is generated in primary and secondary auditory cortices. Second, clicktrain stimuli produce an auditory steady-state response (ASSR), in which neural populations in primary auditory regions are entrained to the modulation frequency for the duration of the clicktrain [19]. In adults, the entrainment in primary auditory cortex is greatest for clicktrains modulated at 40Hz [20]. Measures of inter-trial coherence (ITC) can also be used to measure the ASSR, by quantifying the degree of phase consistency across trials [21]. One advantage of ASSRs is their high test re-test reliability which approaches an intraclass correlation of 0.96, even with a relatively small number of trials [22,23]. Furthermore, ASSRs are modulated by neural development, increasing in power by approximately 0.01 ITC value per year, until early adulthood [24,25]. This increase has been linked with the maturation of superficial cortical layers [26,27]. This makes the ASSR an ideal tool for studying auditory function in developmental conditions such as ASD.

Two studies have measured ASSRs in an ASD context, that is, in ASD participants and in the first-degree relatives of people diagnosed with ASD. Wilson and colleagues reported a reduction in left-hemisphere auditory ASSR power in a group of 10 autistic adolescents, using an early 37-channel MEG system [2]. The second study reported reduced ITC in first-degree relatives of people diagnosed with ASD, with maximal reductions at 40Hz across both hemispheres [1]. Reductions in the ASSR could therefore be an ASD-relevant endophenotype. Additionally, the finding of reduced ITC suggests that dysregulated phase dynamics in bilateral primary auditory cortex could underlie reductions in the ASSR in ASD. However, measures of ITC have not been applied to study the ASSR directly in a group of autistic participants. Additionally, it remains unclear whether reductions in ASSRs are bilateral [1] or unilateral [2] in nature.

As discussed above, auditory stimuli also elicit a more broadband, transient gamma-band response (tGBR) within 0.1s post-stimulus onset [18]. Previously, using auditory clicktrains, Rojas and colleagues [1] reported equivalent tGBRs between the first-degree relatives of people diagnosed with ASD, and controls. However, using sinusoidal auditory tones, several studies have found reduced tGBRs in ASD [8,28,29]. It therefore remains unclear whether both early evoked *and* later sustained gamma-band activity are dysregulated in ASD (also see Kessler, Seymour & Rippon [7], for a review). We therefore opted to analyse tGBRs alongside ASSRs using clicktrain stimuli in a group of autistic participants. However, the primary focus for this study was on the sustained 40Hz response, given the ASD-related differences, previously reported using clicktrain stimuli [1,2].

This study attempted to replicate and extend previous findings showing differences in ASSRs and tGBRs in autism [1,2], within an adolescent population (aged 14-20). We focussed on this age range because adolescence is a crucial period for brain maturation [30,31] and ASSR power increases with age [24,25]. Therefore, we reasoned that ASD-related differences in the ASSR would be more pronounced for an adolescent versus adult population. We also opted to recruit adolescents rather than children for this study, given fixed (adult) size of most MEG helmets and higher levels of compliance in adolescent populations. Data were collected from a group of 18 ASD participants and 18 typically developing controls using a 306-channel MEG system (Elekta Neuromag). An auditory clicktrain stimulus was presented binaurally to participants, to elicit bilateral ASSRs at 40Hz. To investigate prolonged neural entrainment, clicktrain stimuli were presented for a total of 1.5, rather than 0.5s as in previous studies [1,2]. ASSRs were analysed over time, in order to investigate transient changes in 40Hz power and inter-trial coherence. It was hypothesised that, compared with the control group, the ASD group would show reduced ASSR power and ITC at 40Hz for the duration of clicktrain presentation [1,2]. In contrast, it was hypothesised that tGBRs would be equivalent between groups, as previously reported by Rojas & colleagues [1].

## Methods

### Participants

Data were collected from 18 participants diagnosed with ASD and 18 age-matched typically developing controls, see Table 1. ASD participants had a confirmed clinical diagnosis of ASD or Asperger’s syndrome from a paediatric psychiatrist. Participants were excluded from participating if they were taking psychiatric medication or reported epileptic symptoms. Control participants were excluded from participating if a sibling or parent was diagnosed with ASD. MEG data from a further 9 participants was collected but excluded, due to: intolerance to MEG resulting in experimental attrition (2 ASD); movement over 0.5cm (2 ASD, 2 control, see MEG Acquisition); metal artefacts (1 ASD, 1 control); AQ score over 30 (1 control). The movement and AQ thresholds were defined before data collection began.

**Table 1:** Participant demographic and behavioural data. SD = Standard Deviation. * = behavioural scores significantly greater in ASD>control group, t-test, p<.05.

|  | <u>N</u> | <u>Age</u> | <u>Male/Female</u> | <u>Autism Quotient</u><br><u>(Adult) /50</u> | <u>Raven</u><br><u>Matrices</u><br><u>Score /60</u> | <u>Glasgow</u><br><u>Sensory Score</u><br><u>/168</u> | <u>Mind in the</u><br><u>Eyes Score /36</u> |
| --- | --- | --- | --- | --- | --- | --- | --- |
| **ASD** | 18 | Mean = 16.67;<br>SD = 3.2;<br>Range = 14-20 | 14 M; 4 F | Mean = 32.60*;<br>SD = 6.64;<br>Range = 21-46 | Mean = 43.84;<br>SD = 7.93;<br>Range = 27-56 | Mean = 65.33*;<br>SD = 27.69;<br>Range = 27-126 | Mean = 21.88;<br>SD = 4.87;<br>Range = 12-30 |
| **Control** | 18 | Mean = 16.89;<br>SD = 2.8;<br>Range = 14-20 | 15 M; 3 F | Mean = 10.91;<br>SD = 5.43;<br>Range = 6-21 | Mean = 48.71;<br>SD = 5.78;<br>Range = 37-56 | Mean = 38.70;<br>SD = 6.88;<br>Range = 29-50 | Mean = 25.44;<br>SD = 4.03;<br>Range = 17-33 |

### Behavioural Assessments

The severity of autistic traits was assessed using the Autism Quotient (AQ) [32] and sensory traits using the Glasgow Sensory Questionnaire (GSQ) [33]. Using an independent samples t-test, it was shown that both AQ scores, t(34) = 9.869, p<.001 and GSQ scores, t(34) = 3.533, p=.001, were significantly higher in the ASD group, compared with the control group (see Table 1).

Breaking down the GSQ scores further, we observed that our ASD participants showed a heterogeneous pattern of sensory symptoms, with mixtures of hypo- and hyper-sensitivities, across sensory domains (see Supporting Figure 4). Interestingly, auditory scores were the highest amongst any sensory modality, with a mean of 13.9/24. This means that the ASD participants, on average, answered between “Sometimes” and “Often”, when reporting atypical auditory processing on the GSQ.

General non-verbal intelligence was assessed using the Raven’s Matrices Task [34]. Using an independent samples t-test, it was shown that there were no significant group differences in the Raven Matrices Score, t(34) = −1.372, p=.179.

Participants also completed the Mind in the Eyes test [35], however there were no group differences for this test, t(34) = −1.615, p=.116. The Mind in the Eyes test has been recently criticised for measuring emotion recognition rather than an autism-specific deficit in mental state attribution [36], and therefore these scores were not used to investigate correlations between brain patterns and questionnaire measures.

### Paradigm

Whilst undergoing MEG, participants performed an engaging sensory task. Each trial started with a randomised fixation period (1.5, 2.5 or 3.5s), followed by the presentation of a visual grating or auditory binaural click train stimulus. The visual and auditory stimuli were presented randomly, rather than in separate experimental blocks. Only the auditory clicktrain data will be described in this article (please see Seymour et al., 2019 [13] for analysis of the visual grating data). The auditory clicktrain was created from auditory square wave clicks, each of 2ms duration delivered every 25ms for a total of 1.5s. Clicktrains were presented at 80dB (verified using a decibel meter after pneumatic transduction and transmission) binaurally through Etymotic MEG-compatible ear tubes. To keep participants engaged with the task, cartoon pictures of aliens or astronauts were presented after the auditory clicktrain, for a maximum of 0.5s. Participants were instructed to press a response-pad as soon as they were presented with a picture of an alien, but not if they were presented with a picture of an astronaut (maximum response duration allowed was 1.0s). Correct versus incorrect responses were conveyed through 0.5s-long audio-visual feedback (correct: green box, high auditory tone; incorrect responses: red box, low auditory tone). Prior to MEG acquisition, the nature of the task was fully explained to participants and several practice trials were performed. MEG recordings lasted 12-13 minutes and included 64 trials with auditory clicktrain stimuli. Accuracy of picture classification was above 95% for all participants.

**Figure 1.**
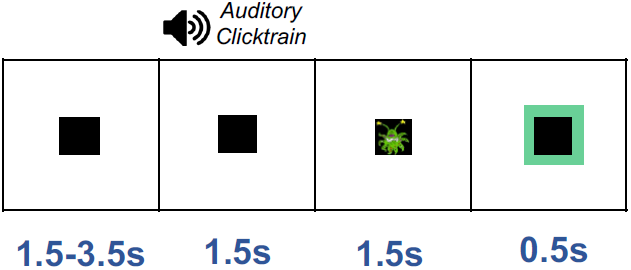
Experimental procedure. (A) Participants performed an audiovisual task, consisting of 1.5–3.5 s baseline period followed by presentation of an auditory clicktrain stimulus for a duration of 1.5s. After this, participants were presented with a cartoon alien or astronaut picture and instructed to only respond when an alien was presented (response time up to 1.5 s), followed by a green or a red framed box for a correct or an incorrect response, respectively. The alien/astronaut stimuli were to maintain attention and do not form part of the analysed data.

### MEG Acquisition

MEG data were acquired using a 306-channel Neuromag MEG scanner (Vectorview, Elekta, Finland) made up of 102 triplets of two orthogonal first-order planar gradiometers and one magnetometer. Acquisition was conducted in a MaxShield™ magnetically shielded room using MaxShield’s™ patented “four layers in one shell” construction. All data were recorded at a sampling rate of 1000 Hz, using default on-line filters between 0.1 - 330 Hz. Internal Active Shielding (Max Shield) was turned off for the recording due to numerous artefacts generated by this technology. Five head position indicator (HPI) coils were applied for continuous head position tracking, and visualised post-acquisition using an in-house Matlab script. Any participant who moved more than a conservative threshold of 5mm in any one direction (x, y or z) were excluded from subsequent analysis. For MEG-MRI coregistration purposes, the locations of three anatomical landmarks (nasion, left and right pre-auricular points), the locations of the HPI coils and 300-500 points from the head surface were acquired using a Polhemus Fastrak digitizer.

### Structural MRI

A structural T1 brain scan was acquired for source reconstruction using a Siemens MAGNETOM Trio 3T scanner with a 32-channel head coil (TE=2.18ms, TR=2300ms, TI=1100ms, flip angle=9°, 192 or 208 slices depending on head size, voxel-size = 0.8×0.8×0.8cm).

### MEG-MRI Coregistration and Cortical Mesh Construction

MEG data were co-registered with participants’ structural MRIs by matching the digitised head-shape data with surface data from the structural scan [37]. Two control participants did not complete a T1 structural MRI and therefore the digitised head-shape data was matched with a database of 95 structural MRIs from the Human Connectome Database [38], using an Iterative Closest Points (ICP) algorithm. The headshape-MRI pair with the lowest ICP error was then used as a ‘pseudo-MRI’ for subsequent steps. This procedure has been shown to improve source localisation performance, in situations where a subject-specific anatomic MRI is not available [39,40]. The aligned MRI-MEG images were used to create a forward model based on a single-shell description of the inner surface of the skull [41] (3000 vertices), using the segmentation function in SPM8 [42]. The cortical mantle was then extracted to create a cortical mesh, using Freesurfer v5.3 [43], and registered to a standard *fs_LR* mesh, based on the Conte69 brain [44], using an interpolation algorithm from the Human Connectome Project [45] (also see: https://goo.gl/3HYA3L). Finally, the mesh was downsampled to 4002 vertices per hemisphere.

### MEG Pre-Processing

Following data inspection, four MEG channels containing large amounts of non-physiological noise (low-frequency drift) were removed from the data, and not included in any of the subsequent pre-processing steps. No channel interpolation was performed. MEG data were pre-processed using Maxfilter (temporal signal space separation, default settings with correlation limit raised to 0.9), which attenuates external sources of noise from outside the head [46]. Further pre-processing steps were performed in Matlab 2014b using the Fieldtrip toolbox v20161024 [47]. Firstly, for each participant the entire recording was band-pass filtered between 0.5-250Hz (Butterworth filter, low-pass order 4, high-pass order 3) and band-stop filtered (49.5-50.5Hz; 99.5-100.5Hz) to remove residual 50Hz power-line contamination and its harmonic. Data were epoched into segments of 4s (1.5s pre, 1.5s post stimulus onset, with 0.5s of padding either side) and each trial was demeaned and detrended. Trials were inspected if the trial-by-channel (magnetometer) variance exceeded 8×10^−23^ (threshold determined from pilot testing), and those containing artefacts (SQUID jumps, eye-blinks, head movement, muscle) were removed. This resulted in the rejection, on average, of 3.4 trials per participant (mean number of trials across participants = 60.6, minimum = 54, maximum = 64). The remaining mean number of trials across participants used for analysis was therefore 60.6 (minimum = 54, maximum = 64). Finally, data were resampled to 200Hz to aid computation time.

### Source-Level Spectral Power

Source analysis was conducted using a linearly constrained minimum variance beamformer [48], which applies a spatial filter to the MEG data at each vertex of the cortical mesh. Only data from orthogonal first-order planar gradiometers were used for source analysis due to current controversy over combining both magnetometers and gradiometers (which have different levels of noise). A covariance matrix for each participant was constructed from the non-averaged, filtered (see below) data, rather than trial-averaged data (as sensor-level data will be made more ‘correlated’ by averaging over trials). Beamformer weights were calculated by combining the gradiometer covariance matrix with leadfield information, with data pooled across baseline and clicktrain periods (−1.5s to 1.5s). Based on recommendations for optimisation of MEG beamforming [49], a regularisation parameter of lambda 5% was applied, due to the rank-deficiency of the data following the Maxfilter procedure.

Whilst the tGBR and ASSR originate from primary auditory cortex, both responses have different frequency ranges and underlying neural generators [25]. Therefore we opted to use separate spatial filters, rather than single spatial filter based on the M100 as used in previous studies [2,50,51]. This decision was based on recent work suggesting that beamformer weights should be optimised for specific data of interest [52].

To localise the ASSR, data were band-pass filtered (Butterworth filter, fifth order) between 35-45Hz. A period of 0.0-1.5s following stimulus onset was compared with a 1.5s baseline period (1.5 to 0.0s before clicktrain onset, also see Figure 2). To localise the tGBR, data were band-pass filtered between 30-60Hz, and a period of 0.0-0.1s following clicktrain onset was compared with a 0.1s baseline period (see Figure 2).

**Figure 2:**
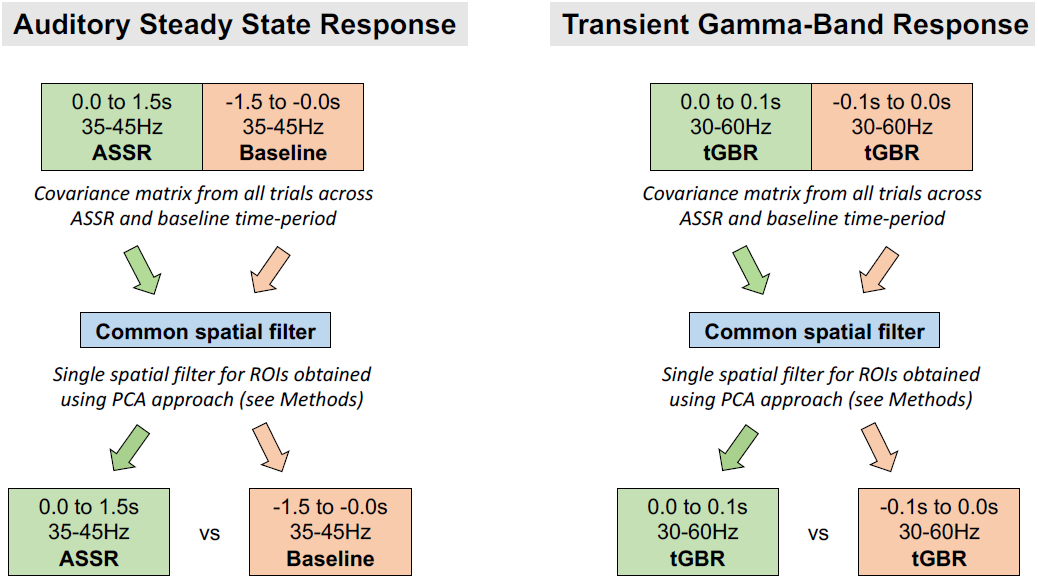
Procedure for Source Analysis. For ASSR beamforming, a common spatial filter was computed using data pooled across ASSR and baseline data. This common filter was then used to localise ASSR/baseline data separately. This process was repeated for tGBR data, but the common spatial filter was computed using a different time and frequency band of interest. ASSR = Auditory Steady State Response. tGBR = Transient Gamma-Band Response.

### ROI definition

Regions of interest (ROI) were selected in bilateral primary auditory (A1) cortices to investigate ASSRs and tGBRs in greater detail. ROIs were defined using a multi-modal parcellation from the Human Connectome Project (Supporting Figure 1, [53]). To obtain a single spatial filter for each ROI (right A1 and left A1 separately), we performed principal components analysis on the concatenated filters of each ROI, multiplied by the sensor-level covariance matrix, and extracted the first component, see [54]. Broadband (0.5-250Hz) sensor-level data were multiplied by this spatial filter to obtain “virtual electrodes”.

### A1 Spectral Power

A1 gamma power (ASSR, tGBR) was analysed using the multi-taper method, as implemented in the Fieldtrip toolbox [47], using discrete prolate spheroidal sequences (Slepian functions). This has been shown to offer an optimal trade-off between time and frequency resolution, and is preferred to Morlet wavelets for high-frequency gamma-band activity [55,56]. Oscillatory power was calculated from 35-45Hz using a 0.5s sliding window (step size 0.02s) with ±5Hz frequency smoothing. Due to the narrow frequency range under investigation, power values were averaged between 35-45Hz. Finally, the percentage change in ASSR power was calculated, using the baseline time-period, i.e. 1.5s before clicktrain presentation. For tGBR power, power values were averaged between 30-60Hz and across time (0-0.1s versus a baseline window 0.1s before clicktrain presentation). From this, the percentage change in tGBR power was calculated.

### A1 Inter-trial Coherence

To assess band-limited phase consistency across trials, we calculated inter-trial coherence (ITC). An ITC value of 0, indicates complete absence of phase consistency, whereas a value of 1 indicates perfect phase consistency across trials [21]. ITC values were converted to Z-values, as recommended by Maris & colleagues [57], to ensure a normal distribution for the statistical analysis (see below).

### Statistical Analysis

For MEG data, statistical analysis was performed using cluster-based permutation tests as implemented in the Fieldtrip toolbox, which have been shown to adequately control the type-I error rate for electrophysiological data [58]. Cluster permutation tests consist of two parts: first an uncorrected independent t-test is performed (two-tailed), and all values exceeding a 5% significance threshold are grouped into clusters. The maximum t-value within each cluster is carried forward. Second, a null distribution is obtained by randomising the data labels (e.g. ASD/control and time) 10,000 times and calculating the largest cluster-level t-value for each permutation. The maximum t-value within each original cluster is then compared against this null distribution, with values exceeding a 5% significance threshold (corrected across both tails, i.e. p<.025 for each tail) deemed significant. Given that only two other MEG studies have used auditory clicktrains to study ASD-related differences [1,2], we adopted a conservative statistical approach and used two-tailed tests in all instances.

For both left and right A1, the following within-group planned statistical contrasts were performed: ASSR power (0.0s to 1.5s) versus baseline (−1.5s to 0.0s); ASSR ITC (0.0s to 1.5s) versus baseline (−1.5s to 0.0s). In addition, the following between-group planned statistical contrasts were performed: control versus ASD ASSR power (0-1.5s post clicktrain onset); control versus ASD ITC (0-1.5s post clicktrain onset); control versus ASD tGBR power (0.0s to 0.1s).

## Results

### ASSR – Power

Whilst ASSRs are known to originate from bilateral primary auditory cortex [19,59], in order to confirm successful source localisation with our pipeline, ASSR power (35-45Hz) was localised on a cortical mesh, using an LCMV beamformer, see Methods. We then calculated the percentage change in 35-45Hz power between 0.0-1.5s post-clicktrain onset versus a 1.5s baseline period (−1.5 to 0.0). As expected, the control group showed maximal increases in power for regions overlapping bilateral primary auditory cortex (Figure 3A) [20,23]. For the ASD group, there were increases in ASSR power for right, but not left, auditory regions, albeit with lower average values than controls (Figure 3B). For an alternative visualisation of results featuring un-thresholded whole-brain statistical maps, see Supporting Information, Figure 3.

**Figure 3.**
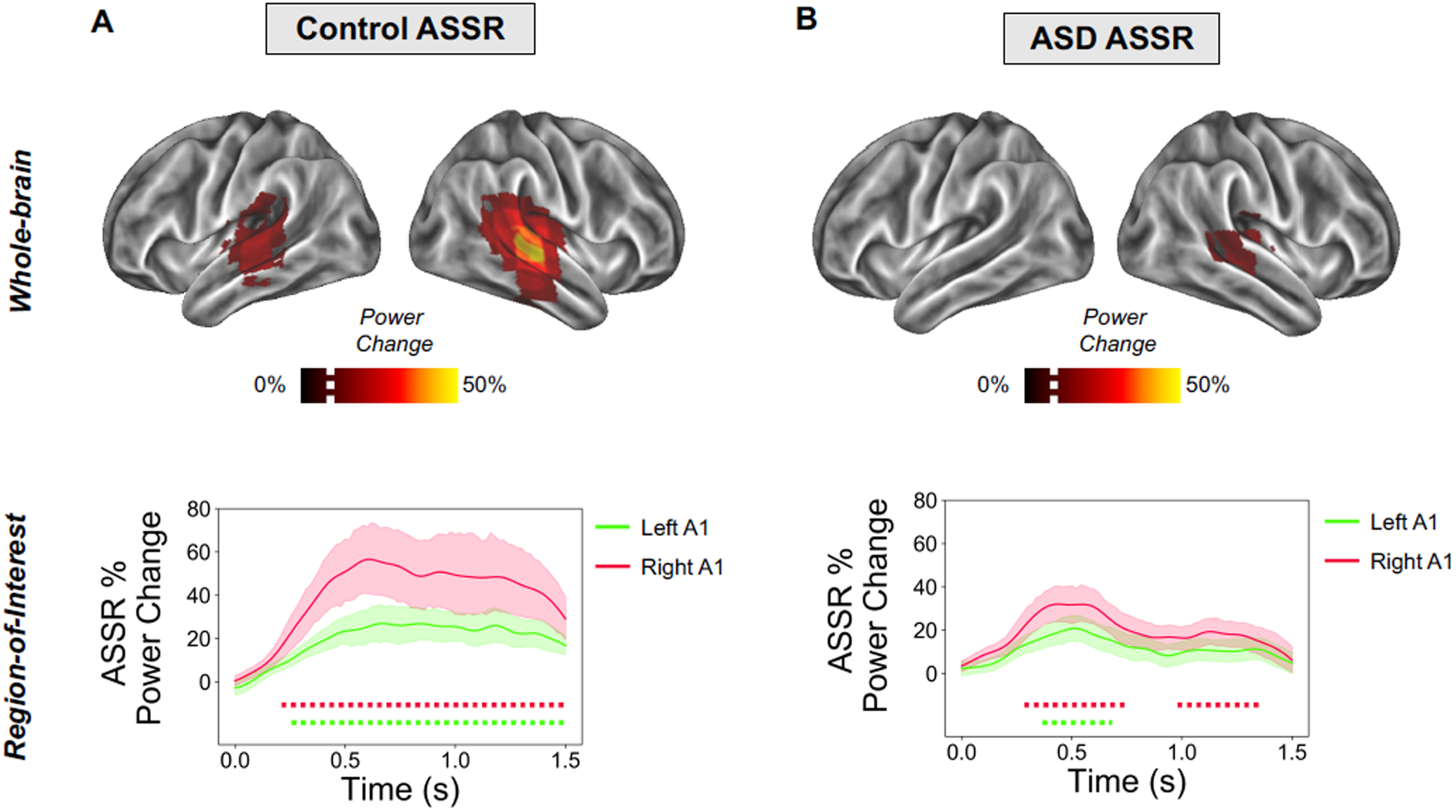
ASSR Power Analysis. Top panels (A, B): ASSR beamformer localization. The percentage change in ASSR power (35-45Hz) is presented on a 3D cortical mesh, thresholded at values greater than 10% (white dotted line on colour scale) for illustrative purposes (for unthresholded images, see Supporting Figure 3), separately for control (A) and ASD (B) groups. Bottom panels (A,B): ASSR in Regions of Interest (ROIs). ROIs were defined in left and right A1 (see Supporting Figure 1) and ASSR oscillatory power was calculated between 35-45Hz. The time-period 0-1.5s post-clicktrain onset was statistically compared with a 1.5s baseline period. Data are plotted separately for: (A) the control group and (B) the ASD group. Dotted lines under the graph indicate times passing a p<.05 threshold (two-tailed) compared to baseline, with different colours corresponding to right A1 (red) and left A1 (green). ASSR = Auditory Steady State Response.

The use of beamforming for bilateral auditory responses has been questioned, due to the potential for mis-localisations resulting from correlated neural sources [48]. However, as noted by Van Veen & colleagues [48] and later by Sekihara & colleagues [60], complete suppression of brain activity, only occurs when the cross-correlation of sources exceeds 0.9. When realistic sources of noise are added to simulated MEG data, complete suppression does not occur [61]. Instead Qurana & Cheyne [61], have shown that for correlated sources at realistic signal-to-noise ratios, beamformers produce a single localisation directly in-between the two sources. Given the clear separation between bilateral auditory sources in our data, as shown in Figure 3, we argue that systematic mis-localisation is unlikely to have occurred. Furthermore, following source analysis we used the online Neurosynth and Neurovault tools to compute the spatial correlation between unthresholded group-level, whole-brain images (see Figure 3; Supporting Figure 3) and several “concept-based meta-analysis maps”, generated from over 10,000 neuroimaging studies [62]. Results, reported in Supporting Table 1, showed the highest correlation with the term “auditory” for both the control (r = .635) and ASD group (r = .471).

ROIs were defined in bilateral auditory cortex (see Supporting Figure 1), to investigate time-frequency responses in greater detail. Oscillatory power was calculated in steps of 0.02s using the multitaper method, and post-stimulus periods (0 to 1.5s) were statistically compared to baseline periods (−1.5 to 0s). Control participants showed increased 35-45z power from 0.24-1.5s for left A1 and 0.21-1.5s for right A1 (Figure 3A bottom panel, times passing a p<.05, two-tailed, threshold are indicated with a dotted line). In contrast, the ASD group showed increased 35-45Hz power for much shorter time windows: in right A1 between 0.31-0.72s and 0.97-1.35s; and for left A1 between 0.41s-0.63s (Figure 3B top panel, times passing a p<.05, two-tailed, threshold are indicated with a dotted line). Next, we statistically compared ASSR 35-45Hz power between groups, for both ROIs. It was found that the control group had greater 35-45Hz power in both right A1, 0.50-1.5s (Figure 4A) and left A1, 0.59 to 1.5s (Figure 4B), compared with the ASD group.

**Figure 4.**
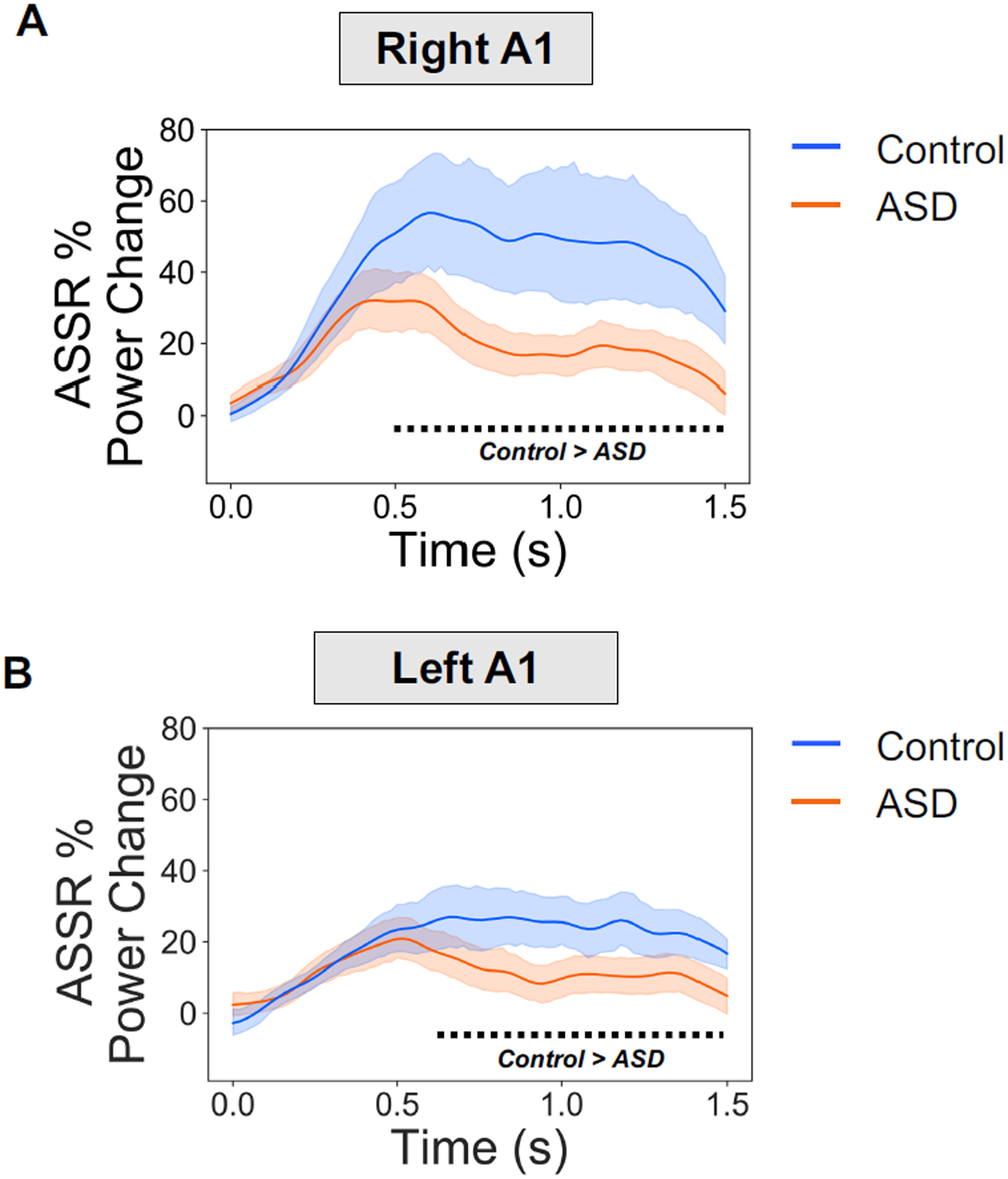
ASSR Power Between Groups. ASSR 35-45Hz power was statistically compared between groups for (A) right A1 and (B) left A1. It was found that the control group had greater ASSR power than the ASD group from 0.52-1.5s in right A1 and 0.60-1.5s for left A1. The black dotted line under the graph indicates times passing a p<.05 threshold (two-tailed) for the Control>ASD contrast.

### ASSR – Intertrial Coherence (ITC)

Next, inter-trial coherence (ITC) was calculated for the A1 ROIs, using the same time-frequency approach as for power. ITC values were Z-scored for statistical analysis [57]. First, we statistically compared the post-clicktrain time-period (0 to 1.5s) with the baseline time-period (−1.5 to 0s). The control group showed statistically significant, p<.05, increases in ITC from 0.1-1.48s for left A1, and 0.14-1.39s for right A1 (Figure 5A). The ASD group showed statistically significant, p<.05, increases in ITC from 0.18-1.50s for left A1, and between 0.12-0.60s, 0.80-0.90, and 1.08-1.5s for right A1 (Figure 5B).

**Figure 5.**
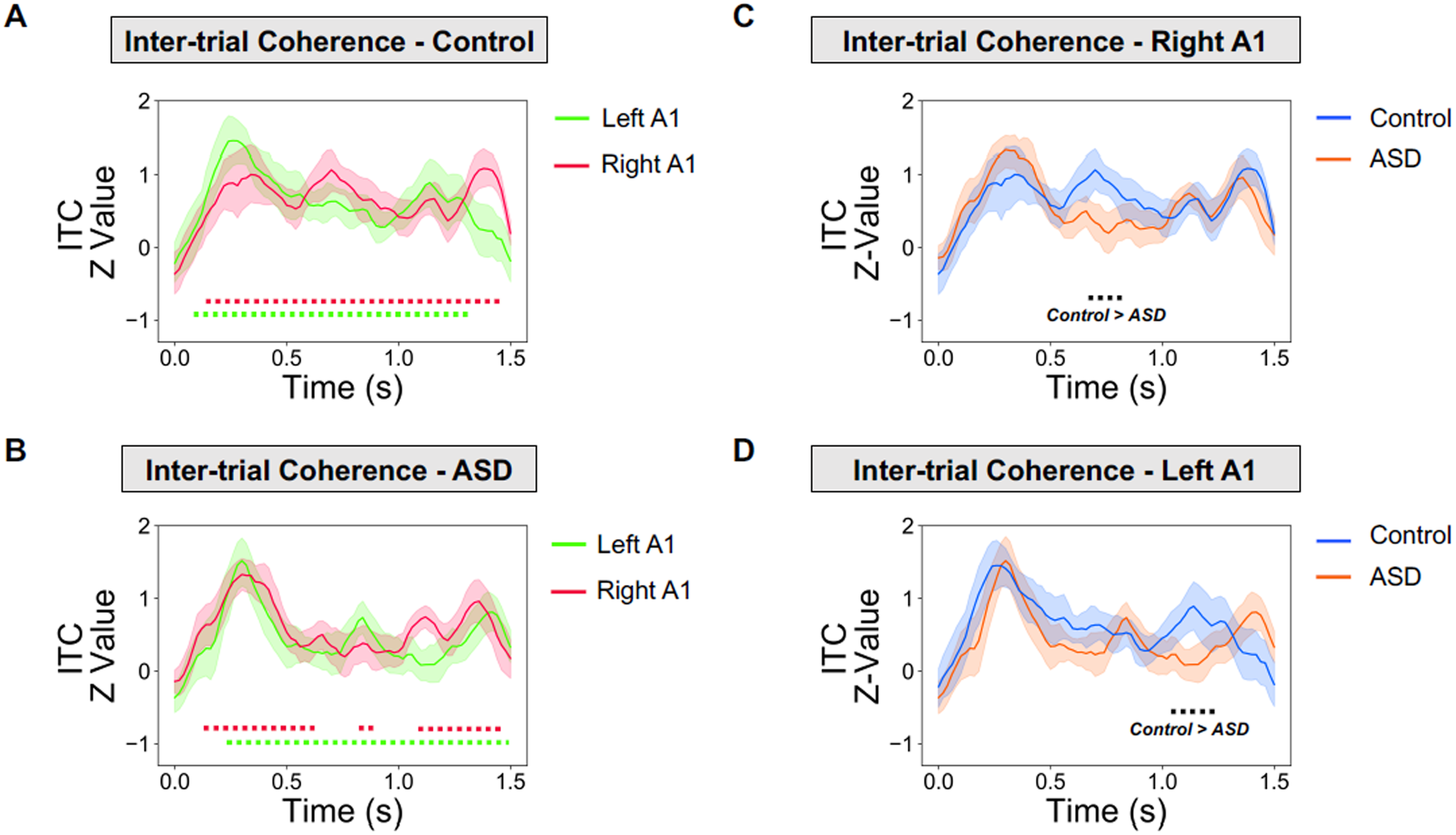
ASSR ITC Analysis. (A-B) 35-45HZ ASSR inter-trial coherence (ITC) results. The time-period 0-1.5s post-clicktrain onset was statistically compared with a 1.5s baseline period. Data are plotted separately for: (A) the control group and (B) the ASD group. Dotted lines under the graph indicate times passing a p<.05 threshold (two-tailed) compared to baseline, with different colours corresponding to right A1 (red) and left A1 (green). (C-D) Between Group 35-45Hz ASSR ITC results. It was found that the control group had greater ASSR ITC than the ASD group from 0.64-0.82s in right A1 (C) and 1.04-1.22s for left A1 (D). The black dotted line under the graph indicates times passing a p<.05 threshold (two-tailed) for the Control (blue) > ASD (red) contrast.

Statistical comparison of ITC between groups showed that the control group had higher ITC in both right A1 (Figure 5C, p<.05) and left A1 (Figure 5D, p<.05), but only within short time-windows from 0.64-0.82s (right A1, Figure 5C) and 1.04-1.22s (left A1, Figure 5D).

### ASSR – Behavioural Data

Next, we investigated whether ASSR responses in the ASD group were correlated with behavioural questionnaire data collected from participants. ASSR power and ITC values were averaged, separately, over those times showing a significant difference (p<.05) between the control and ASD group, as reported in the previous sections (also, see Figure 4 and 5C-D, black dotted lines). Furthermore, given the similar time-course of group differences across right and left A1, ASSR power values were averaged across the ROIs. This was not the case for ITC Z-values, however, which were not averaged across right and left A1. These data were correlated with Autism Quotient (AQ) and Glasgow Sensory Questionnaire (GSQ) scores, for the ASD group, only. There were no significant correlations for ASSR power (Figure 6A, top (AQ) r = .01, p = .96; bottom (GSQ) r = -.17, p = .50), or ITC Z-values (Figure 6B, top (AQ, left A1) r = -.36, p = .14; top (AQ, right A1) r = .11, p = .65; bottom (GSQ, left A1) r = -.35, p = .14; bottom (GSQ, right A1) r = .02, p = .92). The correlation analysis, was repeated for Glasgow Sensory Questionnaire (GSQ) Scores, summed across the six auditory questions only, however no significant correlations were found (p>.05, see Supporting Figure 5).

**Figure 6.**
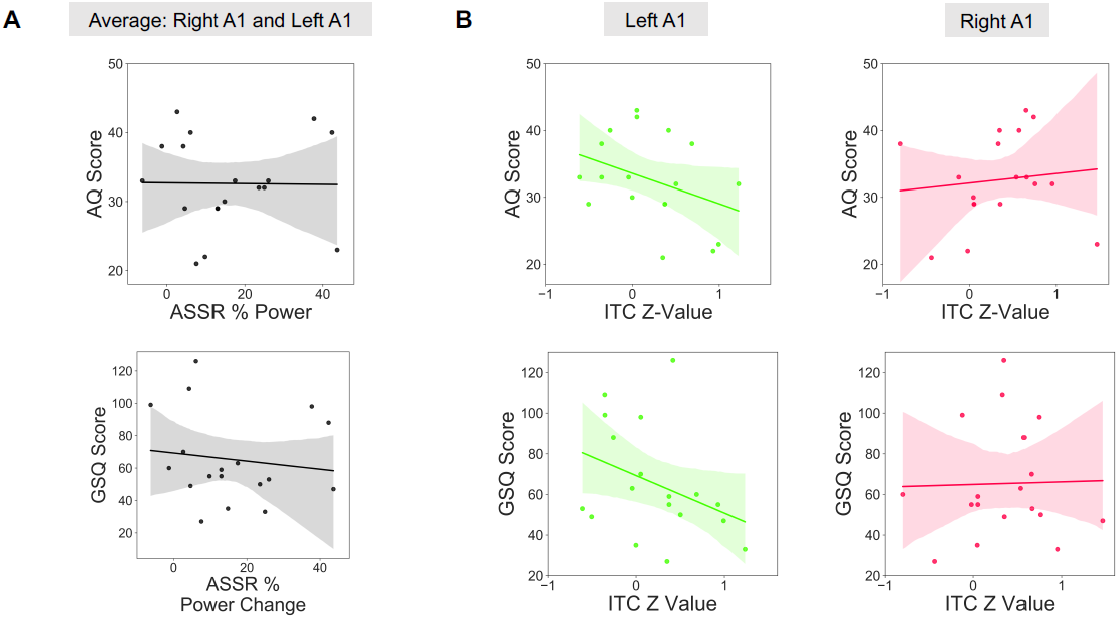
ASSR – Behaviour relationships. Scatter plots to show the relationship between ASSR power, averaged across left/right A1 (A), or ITC Z-values (B), with Autism Quotient (AQ) and Glasgow Sensory Questionnaire (GSQ) Scores. There were no significant (p>.05) correlations for any brain-behaviour relationship. The shaded region indicates 95% confidence intervals. ITC = Intertrial Coherence; ASSR = Auditory Steady State Response.

### tGBR – Source-Level

Transient gamma-band responses to the auditory clicktrain were localised using a beamforming approach (see Methods). As for the ASSR analysis, we first confirmed that the cortical generator(s) of the ASSR originated in bilateral auditory cortex. We calculated the percentage change in 30-60Hz power from 0.0-0.1s post-clicktrain onset compared with a 0.1 baseline period [58]. As expected, both groups group showed maximal increases in tGBR power for regions overlapping with bilateral primary auditory cortex (Figure 7A) [20,23].

**Figure 7.**
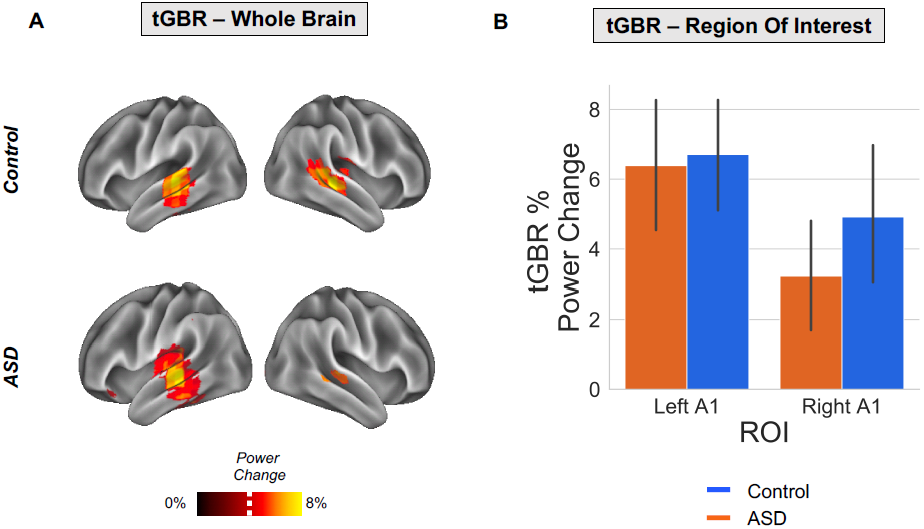
tGBR Analysis. (A) The percentage change in transient gamma-band response, tGBR, power (30-60Hz) is presented on a 3D cortical mesh, thresholded at t > 3.6% (white dotted line) for illustrative purposes, separately for control (top) and ASD (bottom) groups. (B) ROIs were defined in left and right A1 (see Supporting Figure 1). The percentage change in tGBR was plotted separately across ROIs (ASD: blue bar; controls: orange bar). Solid black lines indicate 95% confidence intervals. There were no significant differences in tGBR between groups (p>.05).

Paralleling the ASSR analysis, ROIs were defined in left and right A1. For each ROI and participant, we calculated the percentage change in tGBR power from 30-60Hz, between 0.0-0.1s post-clicktrain onset and a 0.1s baseline period. Statistically comparing groups, it was found that there were no significant differences, p>.05, in tGBR power (see Figure 7B).

## Discussion

This study examined the oscillatory basis of auditory steady state responses (ASSRs) and transient gamma-band responses (tGBR) in a group of 18 autistic adolescents and 18 typically developing controls. We utilised robust source-localisation methods and analysed auditory responses across time. Compared to the ASSR in the control group, we found reduced ∼40Hz power for the ASD group, for regions of interest defined in the left and right primary auditory cortices. Furthermore, there was reduced inter-trial coherence for the autistic group at 40Hz, suggesting that phase dynamics in A1 were less consistent over time. Our results corroborate the notion that auditory brain responses in autism are locally dysregulated [7], especially during sustained gamma-band entrainment (<0.5s post-stimulus onset).

### Auditory Steady State Responses (ASSR) in Autism

Our results are largely consistent with two previous studies which show reduced ASSRs in autistic adolescents [2] and first-degree relatives of people diagnosed with autism [1]. Whilst our study shows reductions in 40Hz power across *both* hemispheres (Figure 3-4), Wilson & colleagues, observed a selective *left*-hemisphere reduction in power [2]. This might be due to the monaural stimulation approach, used by Wilson & colleagues, producing larger hemispheric asymmetries as compared to binaural auditory stimulation [63]. Future work is clearly needed to clarify hemispheric asymmetries in ASSR power for ASD populations [63].

Our results build on the previous literature in several ways. Firstly, by examining sustained ASSRs from 0-1.5s we found that group differences emerged beyond 0.5s post-stimulus onset (Figures 3-4), suggesting that, when driven at gamma frequencies, A1 becomes increasingly dysregulated in ASD compared to controls in a time-dependent manner. This raises the intriguing possibility that sustained, rather than transient, oscillatory activity at gamma-frequencies is affected in autism, perhaps reflecting synaptic dysfunction and an imbalance between excitatory and inhibitory populations of neurons [16]. To investigate this further, future work could parametrically modulate clicktrain duration, intensity, and variability (e.g. perfect 40 Hz vs 38-42Hz, etc.). Secondly, we also found group differences in inter-trial coherence (ITC), with reductions in the autistic group for two short time periods between 0.5 to 1.2s post-stimulus onset (Figure 6). Importantly, measures of ITC are normalised by amplitude and have been shown to be more robust for data with lower signal-to-noise ratios [23]. The reduction in ITC for the autistic group may reflect reduced phase consistency across trials and more idiosyncratic neural responses in autism [64,65], as previously reported for evoked data [66]. However, the reductions in ITC could have also emerged through differences in ASSR power between groups [67]. That said, the time-course of group-differences does diverge between ITC and power (see Supporting Figure 6), with maximum ITC group differences not coinciding with maximum ASSR power differences, which would be expected, if the latter would fully drive the former. In any case, our findings strengthen the claim of reduced ASSRs in autism.

### Transient Gamma-Band Responses in Autism

Unlike ASSRs, there were no group differences in the transient gamma-band (30-60Hz) responses to the clicktrain stimulus (Figure 7). Whilst one previous study using sinusoidal tones reported decreased tGBRs for the first-degree relatives of autistic people, a later study using auditory clicktrains, found no group differences in either power or ITC [1]. More generally, findings of transient/evoked gamma-band power across sensory domains are very mixed, with both increases and decreases reported (reviewed in [7]). The divergence between steady-state and transient gamma in this study has implications for potential oscillopathies in ASD, as differences in gamma power may depend on the time-period under investigation as well as the underlying neural circuits generating gamma oscillations [25].

### ASSRs as Markers of Dysregulated Local Activity

There has been recent interest in characterising atypical patterns of gamma-band oscillations in autism, due to their link with local cortical function and connectivity [7]. The precise E-I mechanisms underlying gamma generation are well characterised, for a review see [14]. Of particular importance is the functional inhibition of pyramidal neurons by fast-spiking interneurons via binding of the neurotransmitter gamma-aminobutyric acid (GABA) [14,68]. Relatedly, there is emerging evidence showing GABA dysfunction in autism [68]. Reduced gamma-band steady-state responses in autism may therefore reflect dysregulated neuronal inhibition, resulting in E-I imbalance [16]. As argued by Kessler, Seymour & Rippon [7], this local dysregulation could result in both hyper and hypo-sensitivities in ASD, depending on the particular sensory input, and the degree of top-down modulatory processes employed by individuals [13]. To quantify the precise mechanisms underlying reduced gamma-band ASSRs, future studies could utilise dynamic causal modelling of A1 neuronal circuits [69], combined with parametric modulations of ASSRs (e.g. duration, frequency) and participant attention [70]. It would also be interesting to use more naturalistic auditory stimuli, for example speech stimuli [71,72], to investigate whether neural entrainment is affected more generally in ASD.

It should also be noted that ASSRs are not simply generated via the linear accumulation of transient evoked responses [20,73,74]. Instead, the ASSR may reflect a sustained non-linear neural response at the input stimulation frequency and its harmonics, peaking at the system’s preferred modulation rate [20]. In support of this, Edgar & colleagues (2016) report that in children, ASSRs are difficult to detect, despite measurable auditory evoked responses [25]. Similarly, our data show intact auditory evoked fields (see Supporting Figure 2) and transient gamma-band responses in autism (Figure 7), in the presence of a reduced ASSR (Figure 4). Rather than a generalised gamma-band dysfunction in autism, our data suggest a more nuanced reduction in the non-linear dynamics underlying steady-state auditory gamma [1]. Interestingly, an MEG study examining somatosensory processing in ASD showed reduced frequency harmonics at 50Hz [12], while Vilidaite and colleagues reported a reduction in harmonic EEG responses during visual steady-state stimulation in autistic adults [75]. Furthermore, two MEG studies revealed reduced alpha-gamma phase-amplitude coupling in the visual system in ASD [14, 18]. Overall, this suggests that non-linear aspects of local cortical processing could be dysregulated across sensory domains in ASD [8].

ASSRs are developmentally relevant, increasing by approximately 0.01 ITC value per year [24,25,50]. This trajectory may reflect the continuing development of superficial layers of cortex where gamma-band oscillations predominantly originate [27]. We hypothesise that the ASD-related reduction in ASSRs reported in this study results from an atypical trajectory of gamma-band maturation, in line with developmental disconnection theories of autism [76]. Given that the 40Hz ASSRs continue to mature throughout late adolescence and adulthood [50], it remains to be established, whether the development of ASSRs in ASD is simply delayed, or whether reductions persist throughout life. To investigate this further, future studies should use high-powered longitudinal ASD samples and age-appropriate MEG systems [77], to characterise ASSR development throughout childhood, adolescence and into adulthood [78]. If confirmed, divergent ASSR trajectories could act as important autism-relevant markers of intervention efficacy [79].

### Limitations

In this study, formal clinical ASD assessment of our participants, e.g. the ADOS [80], was not performed. We therefore implemented strict participant exclusion criteria, only including autistic participants with a confirmed clinical diagnosis of ASD or Asperger’s syndrome. Between groups, there were significant differences in autistic and sensory traits, measured using two self-report questionnaires (Table 1). However, upon closer inspection of behavioural data (see Supporting Figure 5), the ASD group showed a mixture of hyper- and hypo-sensitive traits between different sensory modalities making precise brain-behavioural correlations problematic. This may explain the lack of relationship between ASSR power/ITC and AQ/GSQ scores in ASD (Figure 6). Brain-behaviour relationships might be better quantified using MEG in combination with psychophysical tests of auditory perception and formal clinical assessments.

## Declarations

### Ethics approval and consent to participate

All experimental procedures complied with the Declaration of Helsinki and were approved by the Aston University, Department of Life & Health Sciences ethics committee. Participants and a parent/guardian gave written informed consent before participating in the study.

### Consent for publication

N/A

### Availability of data and materials

The data that support the findings of this study are available on reasonable request from corresponding author, R.S., in a preprocessed and de-anonymized form. The raw data are not publicly available due to ethical restrictions.

### Competing Interests

The authors wish to declare the research was conducted in the absence of any commercial or financial relationships that could be construed as a potential conflict of interest.

### Funding

The Wellcome Trust, Dr Hadwen Trust and Tommy’s Fund supported research-scanning costs. Robert Seymour was supported by a cotutelle PhD studentship from Aston University and Macquarie University. Paul F. Sowman was supported by the Australian Research Council [DP170103148].

### Author Contributions

RS: Study design; data collection; data analysis; data interpretation; manuscript writing.

KK: Study design; data interpretation; manuscript writing.

GR: Study design; data interpretation; manuscript writing.

PFS: Data interpretation; manuscript writing.

GGW: Data collection; data interpretation.

## Acknowledgments

We wish to thank: the volunteers who gave their time to participate in this study; Dr Shu Yau and Dr. Michael Hall for help with MRI data acquisition; and Dr Jon Brock for intellectual contributions to experimental design.

## Supporting Information

**Supporting Figure 1:**
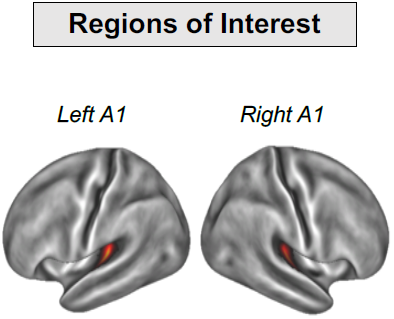
Regions of interest were defined in left and right Primary Auditory Cortex, according to HCP-MMP 1.0 Atlas.

**Supporting Figure 2:**
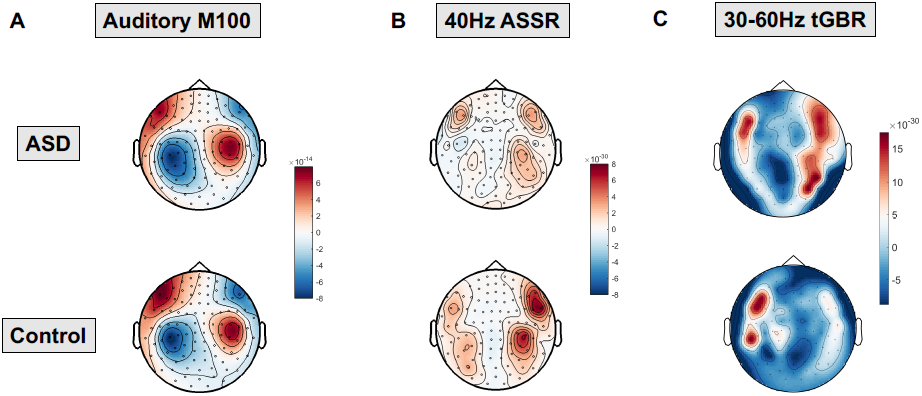
Sensor-level Analysis. **(A)** Group average topo-plot for the auditory M100 event-related field, magnetometers shown. **(B)** Group average topo-plot for auditory steady state responses (ASSR) at 40Hz **(C)** Group average topo-plot of the transient gamma-band response (tGBR), 30-60Hz, 0.0-0.1s. Scales represent MEG field strength, baseline-corrected, with units of Tesla/cm.

**Supporting Figure 3:**
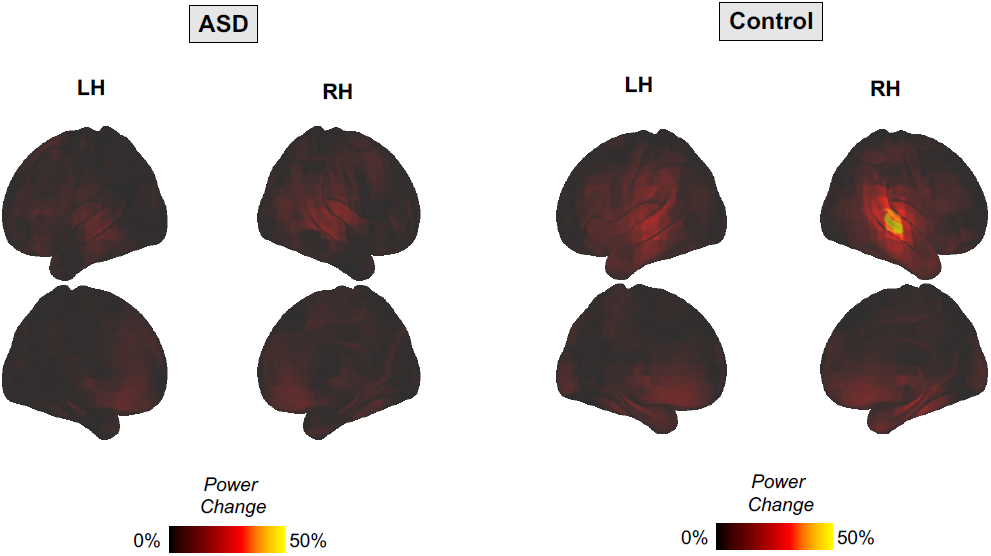
Whole-brain maps showing changes in ASSR power, corresponding to Figure 2 in the main text.

**Supporting Figure 4:**
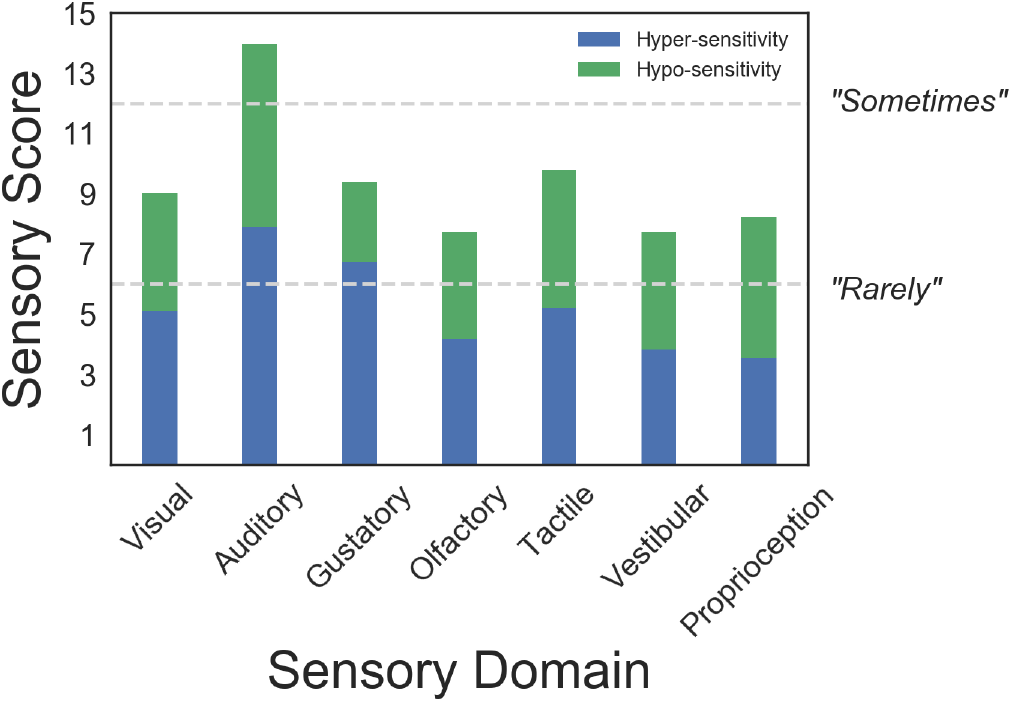
Responses to the Glasgow Sensory Questionnaire were grouped by sensory domain (maximum score = 20) and hypo- / hyper-sensitivity (green and blue bars respectively). Our data show a heterogeneous pattern of sensory symptoms, with mixture of hypo- and hyper-sensitivities. Auditory symptoms scored 13.9/20 corresponding to questionnaire answers closest to “Sometimes”.

**Supporting Figure 5:**
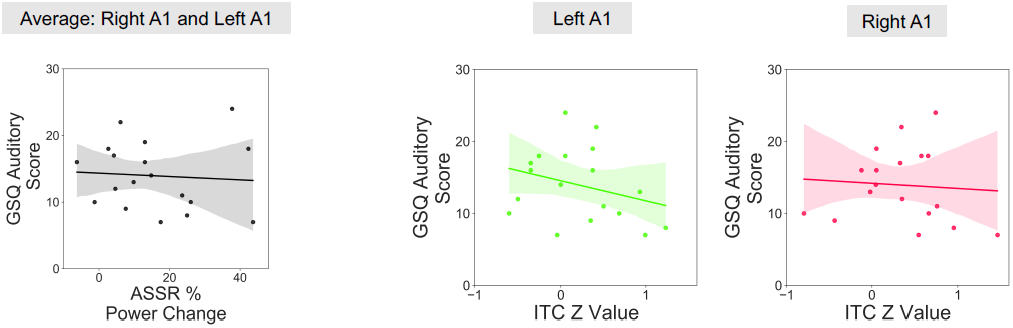
Scatter plots to show the relationship between ASSR power, averaged across left/right A1, and Glasgow Sensory Questionnaire (GSQ) Scores, summed across the six auditory questions only. There were no significant (p>.05) correlations for ASSR power, r = -.07, p=.77, or for ITC Z-Value (left A1), r = -.29, p=.24, (right A1), r = -.07, p = .76. The shaded region indicates 95% confidence intervals. ITC = Intertrial Coherence; ASSR = Auditory Steady State Response.

**Supporting Figure 6:**
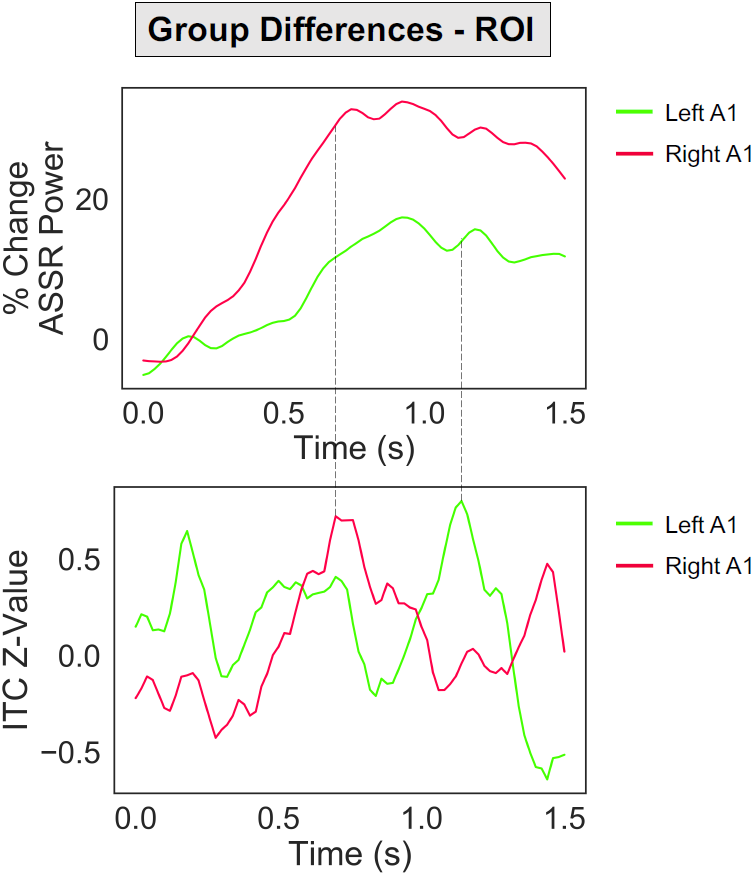
For ASSR power (top) and ITC power (bottom), the group average for the ASD group was subtracted from the group average for controls (at each time bin) and plotted for each ROI (right A1: red; left A1: green). The black dotted lines link the times maximum ITC group differences for left and right A1 with the corresponding data for ASSR power.

**Supporting Table 1:** Results of the meta-analytical decoding (top 10 terms shown). The correlation term corresponds to the r-value between the whole-brain unthresholded map and the Neurosynth concept-based meta-analysis maps, generated from over 10,000 neuroimaging studies.

|  | Control |  | ASD |  |
| --- | --- | --- | --- | --- |
| *Rank* | *Search Term* | *Correlation* | *Search Term* | *Correlation* |
| 1 | auditory | 0.635 | auditory | 0.471 |
| 2 | superior temporal | 0.595 | auditory cortex | 0.427 |
| 3 | auditory cortex | 0.57 | superior temporal | 0.424 |
| 4 | sounds | 0.567 | sounds | 0.408 |
| 5 | listening | 0.557 | sound | 0.404 |
| 6 | sound | 0.552 | listening | 0.398 |
| 7 | temporal | 0.55 | heschl gyrus | 0.392 |
| 8 | planum temporale | 0.539 | heschl | 0.39 |
| 9 | temporale | 0.539 | primary auditory | 0.389 |
| 10 | planum | 0.531 | planum temporale | 0.388 |

## Notes

### Competing Interest Statement

The authors have declared no competing interest.

### Summary of Updates

Minor revisions (June 2020)

